# Collagen co-localised with macrovesicular steatosis for fibrosis progression in non-alcoholic fatty liver disease

**DOI:** 10.1101/2023.02.10.528084

**Authors:** Xiao-Xiao Wang, Rui Jin, Xiao-He Li, Qiang Yang, Xiao Teng, Fang-Fang Liu, Nan Wu, Hui-Ying Rao, Feng Liu

**Author notes:** **Corresponding author:** Feng Liu, MD.

## Abstract

Non-alcoholic fatty liver disease (NAFLD) is a commonly occurring liver disease; however, its exact pathogenesis is not fully understood. The purpose of this study was to quantitatively evaluate the progression of steatosis and fibrosis by examining their distribution, morphology, and co-localisation in NAFLD animal models. qSteatosis showed a good correlation with steatosis grade (*R***: *0.823–0.953***, *P*<0.05) and demonstrated high performance (area under the curve [AUC]: 0.617–1) in all six mouse models. Based on their high correlation with histological scoring, qFibrosis containing four shared parameters were selected to create a linear model that could accurately identify differences among fibrosis stages (AUC: 0.725–1). qFibrosis co-localised with macrosteatosis generally correlated better with histological scoring and had a higher AUC in all six animal models (AUC: 0.846–1). Quantitative assessment using second-harmonic generation/two-photon excitation fluorescence imaging technology can be used to monitor different types of steatoses and fibrosis progression in NAFLD models. The collagen co-localised with macrosteatosis could better differentiate fibrosis progression and might aid in developing a more reliable and translatable fibrosis evaluation tool for animal models of NAFLD.

## Introduction

Non-alcoholic fatty liver disease (NAFLD) is defined as the accumulation of fat in the liver without excessive alcohol consumption or other known liver pathologies. NAFLD encompasses a spectrum including “simple steatosis” (non-alcoholic fatty liver), non-alcoholic steatohepatitis (NASH), fibrosis, and cirrhosis or hepatocellular carcinoma (Golabi et al, 2021; Huang et al., 2021; Paik et al., 2020). NAFLD is emerging as the most common type of chronic liver disease worldwide and might become the main indication for liver transplantation (Terrault and Pageaux, 2018; Younossi et al., 2021). However, the exact pathogenesis and progression of NAFLD are not fully understood (Diehl et al, 2017; Tilg et al., 2021) and NASH still lacks regulatory-approved pharmacotherapy (Ferguson and Finck, 2021; Friedman et al. 2018]. Therefore, it is crucial to design effective interventions that can be safely used to treat patients with NAFLD.

With the limitations in obtaining human samples and performing experimental drug studies in humans, NAFLD animal models are necessary to test the different pathogenic mechanisms and therapeutic targets of human NAFLD (Berardo et al., 2020; Ipsen et al., 2020; Nevzorov et al., 2020; Reimer et al., 2020). Different murine models of NAFLD have been developed using diet, chemical induction, genetic modification, or a combination of these actions. However, at present, no other scoring system has been applied for histological evaluation, except for the Brunt or NASH Clinical Research Network (NASH CRN) scoring system (Brunt et al., 1999; Kleiner et al., 2005). This system is nonlinear and semiquantitative and cannot reflect the minor variability and extent of injury, especially for collagen changes. In addition, subjective evaluations using this system exhibited inter- and intra-observer variabilities. Consequently, precise, objective, and dynamic evaluation of liver histology in preclinical NAFLD animal models is valuable for developing more effective drugs.

The main histological features of NAFLD, including steatosis, inflammation, ballooning, and fibrosis, are often evaluated in preclinical models investigating novel therapies. Among the four features, steatosis is the most important diagnostic feature of NAFLD and is associated with progression to steatohepatitis and fibrosis (Kleiner et al., 2019; McPherson et al., 2015). Liver fibrosis is the most principal prognostic factor in NAFLD and is correlated with future liver-related events and long-term overall mortality (Dulai et al., 2017; Hagström et al., 2017; Vilar-Gomez et al., 2018). Accurate assessment of steatosis and fibrosis severity could contribute to the precision of drug efficacy evaluation, which is crucial for drug development for this disease.

Second-harmonic generation (SHG)/two-photon excitation fluorescence imaging (TPEF) techniques have been reported in the development of new algorithms to assess steatosis and fibrosis in the liver in the past decade (Chang et al., 2018; Liu et al., 2017; Liu et al., 2020; Sun et al., 2008; Wang et al., 2017). It is unclear whether this purely quantitative method can be used in different NAFLD mice models. In addition, unlike macrovesicular steatosis mainly observed in human patients with NAFLD, macrovesicular and microvesicular steatosis often coexist in animal models. Therefore, developing a new objective evaluation system is particularly important for quantifying steatosis and fibrosis progression in preclinical NAFLD animal models. Rodent diets containing 45% kcal fat are very common, which are equally effective in promoting obesity. Fructose, which is recognized to promote hepatic de novo lipogenesis, lipid accumulation, and insulin resistance, typically results in steatohepatitis progressing to moderate fibrosis. Carbon tetrachloride (CCl4) enhances the effects of high-fat diet (HFD) on the rapid development of NASH and fibrosis. Therefore, this study focused on six different mice NAFLD models associated with diet, fructose, and CCl4 induction and developed an automated evaluation system by combining SHG/TPEF microscopy and an adaptive quantification algorithm to distinguish the progression of steatosis and fibrosis.

## Results

### Histology in different NAFLD mouse models

Histological examination (hematoxylin and eosin [H&E] staining) demonstrated that macro-and microvesicular steatosis were clearly visible after 16 weeks in all six NAFLD models. The collagen pattern detected by SHG was highly consistent with that of light microscopic appearance and histochemical staining (Fig. S1 and S2). No signs of stage 2 fibrosis were detected in western diet (WD), WD plus fructose in drinking water (WDF), HFD, or HFG plus fructose in drinking water (HFDF) mice until 24 weeks. However, stage 3 to 4 fibrosis was observed in the WDF+CCl4 and HFDF+CCl4 mice after 16 weeks. CCl4 treatment may accelerate HFD- and WD-induced steatohepatitis and fibrosis.

### Good correlations were observed between the histopathology and SHG/TPEF assessments of steatosis in all mouse models

Liver steatosis progression was quantified by examining the fat vacuoles in the TPEF channel. Overall, 45 steatosis parameters were extracted from the whole-tissue images. Good correlations were observed for all steatosis parameters with steatosis scores in all six NAFLD mouse models (Table 1 and Fig. S3). For representative parameters, such as %Area (percentage of steatosis in the overall region), %MacroArea (percentage of macrosteatosis in the overall region), and %MicroArea (percentage of microsteatosis in the overall region), the Spearman correlations ranged from 0.823 to 0.953 (*P*<0.05) in all mouse models (Table S1). In addition, steatosis area parameters demonstrated high area under the receiver operating characteristic (ROC) curve (AUC) values for differentiating steatosis grades (0.810–1).

**Table 1.** ROC analysis of the representative steatosis parameters in differentiating steatosis grades in the six mouse models.

| Steatosis grade | WD |  |  | WDF |  |  | WDF+CCI4 |  |  |
| --- | --- | --- | --- | --- | --- | --- | --- | --- | --- |
|  | %Area | %MacroArea | %MicroArea | %Area | %MacroArea | %MicroArea | %Area | %MacroArea | %MicroArea |
| 0vs.123 | 1 | 1 | 0.953 | NA | NA | NA | NA | NA | NA |
| 01vs.23 | NA | NA | NA | NA | NA | NA | 0.965 | 0.983 | 0.632 |
| 012vs.3 | 1 | 0.990 | 0.899 | 1 | 1 | 1 | 0.967 | 0.892 | 0.617 |
| Steatosis grade | HFD |  |  | HFDF |  |  | HFDF+CCI4 |  |  |
|  | %Area | %MacroArea | %MicroArea | %Area | %MacroArea | %MicroArea | %Area | %MacroArea | %MicroArea |
| 0vs.123 | NA | NA | NA | NA | NA | NA | 1 | 0.949 | 1 |
| 01vs.23 | NA | NA | NA | 1 | 1 | 0.932 | 1 | 0.921 | 0.905 |
| 012vs.3 | 1 | 1 | 1 | 0.893 | 0.810 | 1 | 1 | 0.857 | 0.964 |

Abbreviations: ROC: receiver-operating-characteristics curve; WD, Western diet; WDF, WD+fructose; CCI4, carbon tetrachloride; HFD, high-fat diet; HFDF, HFD+fructose; NA, not available

### The qFibrosis index combining four shared morphological parameters faithfully recapitulated the fibrosis staging

Based on their quantitative trends of fibrosis stages and systemic AUC analyses, four shared parameters of collagen string (#LongStrPS, #ThinStrPS, #ThinStrPSAgg and #LongStrPSDis) were selected to combine qFibrosis indices, which showed good correlation with fibrosis stages in all animal models (*R*: 0.501–0.911, *P*<0.05) (Fig. S4, Table S2). These indices can also accurately identify differences among fibrosis stages (AUC: 0.725–1) (Table 2).

**Table 2.**
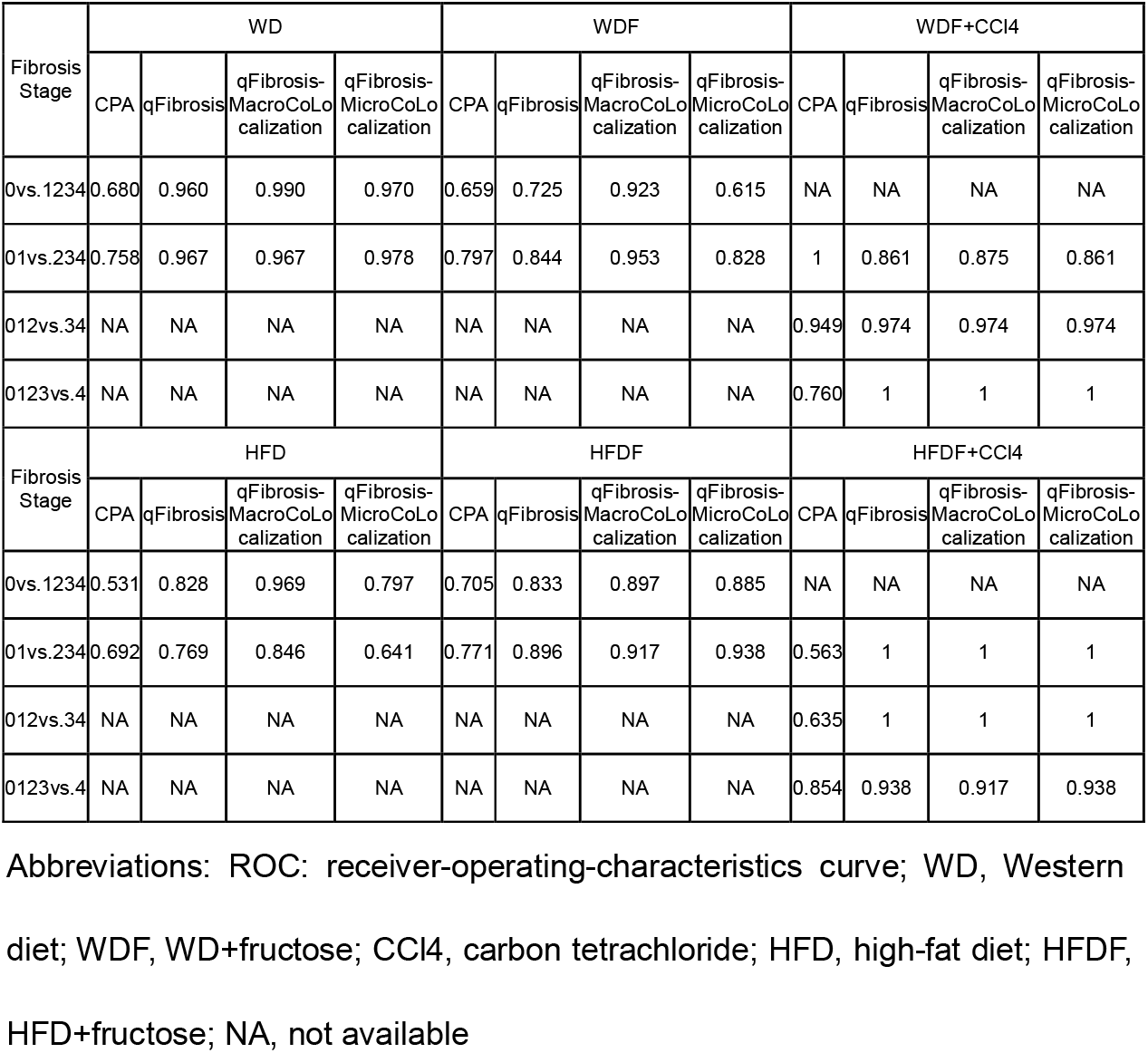
ROC analysis of the representative fibrosis parameters in differentiating fibrosis stages in six mouse models.

| Fibrosis Stage | WD |  |  |  | WDF |  |  |  | WDF+CCl4 |  |  |  |
| --- | --- | --- | --- | --- | --- | --- | --- | --- | --- | --- | --- | --- |
|  | CPA | qFibrosis | qFibrosis-MacroCoLocalization | qFibrosis-MicroCoLocalization | CPA | qFibrosis | qFibrosis-MacroCoLocalization | qFibrosis-MicroCoLocalization | CPA | qFibrosis | qFibrosis-MacroCoLocalization | qFibrosis-MicroCoLocalization |
| 0vs.1234 | 0.680 | 0.960 | 0.990 | 0.970 | 0.659 | 0.725 | 0.923 | 0.615 | NA | NA | NA | NA |
| 01vs.234 | 0.758 | 0.967 | 0.967 | 0.978 | 0.797 | 0.844 | 0.953 | 0.828 | 1 | 0.861 | 0.875 | 0.861 |
| 012vs.34 | NA | NA | NA | NA | NA | NA | NA | NA | 0.949 | 0.974 | 0.974 | 0.974 |
| 0123vs.4 | NA | NA | NA | NA | NA | NA | NA | NA | 0.760 | 1 | 1 | 1 |
| Fibrosis Stage | HFD |  |  |  | HFDF |  |  |  | HFDF+CCl4 |  |  |  |
|  | CPA | qFibrosis | qFibrosis-MacroCoLocalization | qFibrosis-MicroCoLocalization | CPA | qFibrosis | qFibrosis-MacroCoLocalization | qFibrosis-MicroCoLocalization | CPA | qFibrosis | qFibrosis-MacroCoLocalization | qFibrosis-MicroCoLocalization |
| 0vs.1234 | 0.531 | 0.828 | 0.969 | 0.797 | 0.705 | 0.833 | 0.897 | 0.885 | NA | NA | NA | NA |
| 01vs.234 | 0.692 | 0.769 | 0.846 | 0.641 | 0.771 | 0.896 | 0.917 | 0.938 | 0.563 | 1 | 1 | 1 |
| 012vs.34 | NA | NA | NA | NA | NA | NA | NA | NA | 0.635 | 1 | 1 | 1 |
| 0123vs.4 | NA | NA | NA | NA | NA | NA | NA | NA | 0.854 | 0.938 | 0.917 | 0.938 |

Abbreviations: ROC: receiver-operating-characteristics curve; WD, Western diet; WDF, WD+fructose; CCl4, carbon tetrachloride; HFD, high-fat diet; HFDF, HFD+fructose; NA, not available

Furthermore, the performances of qFibrosis indices versus collagen proportional area (CPA) for scoring fibrosis were evaluated using ROC analysis; these AUC values were mostly higher than those using CPA in differentiating fibrosis stages (0.725–1 vs. 0.531–1) (Table 2).

### qFibrosis co-localised with macrosteatosis showed superior performance compared to CPA, qFibrosis, and qFibrosis co-localised with microsteatosis in evaluating fibrosis severity

The relationship between steatosis and fibrosis progression was further analysed by examining the co-localisation (Fig. 1). The results revealed that qFibrosis co-localised with macrosteatosis was generally correlated better with histological scoring than qFibrosis (R: 0.598–0.914, *P*<0.05) and qFibrosis co-localised with microsteatosis (R: 0.516–0.946, *P*<0.05) in most animal models. Furthermore, using ROC analysis, the AUC values of qFibrosis co-localised with macrosteatosis for the detection of different stages were >0.875 (AUC: 0.875–1), whereas the AUC values of qFibrosis were >0.725 (AUC: 0.725–1) and the AUC values of qFibrosis co-localised with microsteatosis were >0.615 (AUC: 0.615–1) (Table 2).

**Figure 1.**
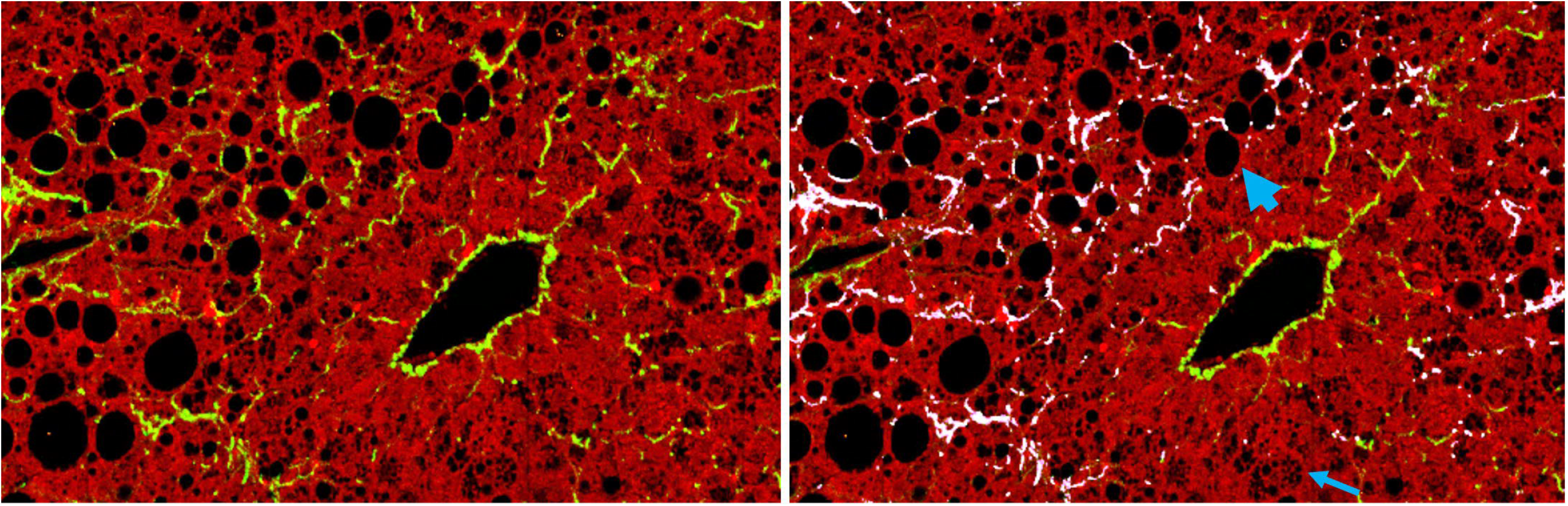
Collagen co-localised with steatosis. The subfigure on the left is a raw ROI image; the subfigure on the right is the corresponding annotated image, where white-coloured collagen fibres are co-localised with steatosis. The bold and thin arrow points to an example of macro- and microsteatosis, respectively.

## Discussion

Animal models remain the best way to entirely study the pathophysiology of liver diseases and develop new treatments. However, few studies have analysed the possibility of automated NAFLD diagnosis using imaging techniques in animal models. In this study, we established an automated evaluation system, qFibrosis co-localised with macrosteatosis, which is superior to the fibrosis staging scores and CPA evaluation in six different NAFLD models.

Hepatic steatosis is the key defining feature of NAFLD. Animal models often combine various degrees of macro- and microsteatosis. Some studies using animal models have analysed new techniques that validate automated computerized systems for NAFLD diagnosis and staging. GE et al. (2010) first evaluated hepatic steatosis based on the quantitative digital analysis of Oil Red O in various mouse models and demonstrated that the area fraction was significantly associated with triglyceride deposition in the liver. Furthermore, Sethunath et al. (2018) analysed macro- and microsteatosis of fatty liver disease in murine models using machine-learning modalities. Ramot et al. (2020) compared the manual semiquantitative microscope-based assessment with data output obtained from artificial intelligence software using a deep learning algorithm and showed an important association (*r*=0.87, *P*<0.01) among the semiquantitative methods performed by a human pathologist and automated diagnostic tools. Our results also showed good correlations between histopathology and SHG/TPEF-based steatosis parameters with steatosis scores in all our mouse models. This suggests that SHG/TPEF can be used to assess hepatic steatosis changes in different NAFLD mouse models.

The fibrosis stage is the main histological predictor of liver-specific outcomes. Accurate and quantitative evaluation of liver fibrosis in NAFLD is very important for predicting the risk of complications and tracking disease progression and should be considered as an effective endpoint in clinical trials of antifibrotic drugs. De Rudder et al. (2020) demonstrated an automated method of measuring the total content of collagen in the picrosirius red-stained liver section regardless of collagen distribution. Roth et al. (2018) demonstrated that INT-767 reduced collagen fibre signals using SHG imaging and suggested that SHG could be used to measure collagen improvements in preclinical NASH mice. Furthermore, based on the trends with respect to fibrosis stages and systemic AUC analyses, we selected qFibrosis by combining four parameters (#LongStrPS, #ThinStrPS, #ThinStrPSAgg, and #LongStrPSDis) to faithfully recapitulate the Ishak scores better than CPA. This suggests that the distribution, length, and width of collagen might be superior to total collagen quantitation and lay the foundation for the assessment liver fibrosis in NAFLD based on SHG microscopy.

In liver tissue with NASH, perisinusoidal fibrosis has a distinctive pattern and is most likely associated with the severity of steatosis (Chalasani et al., 2008). All four parameters for qFibrosis were derived from the perisinusoidal collagen. Therefore, we performed a co-localisation analysis between qFibrosis and steatosis. Our results revealed that qFibrosis co-localised with macrosteatosis generally correlated better with histological scoring than the original qFibrosis and qFibrosis co-localised with microsteatosis in most animal models. Although macrosteatosis is mostly observed in patients with NAFLD, HFD- and WD-treated mice often show progression of macro- and microsteatosis (Ding et al., 2018; Ganz et al., 2014). Transitions and associations exist between macro- and microsteatosis, and it has been hypothesized that large fat droplets are formed by the fusion of small droplets initially found on the surface of the endoplasmic reticulum (Martin and Parton, 2006; Tandra et al., 2011). Quantitative morphometric analyses revealed that INT-767 significantly reduced vesicular lipid droplet size, suggesting that macrovesicular steatosis is positively correlated with the severity of lobular inflammation and fibrosis (Roth et al., 2018). Therefore, qFibrosis co-localised with macrosteatosis could have more discriminative power to precisely reflect the dynamics of fibrosis progression in different NAFLD models.

All these NAFLD models have affirmed that SHG microscopy is an invaluable new platform to study and quantify steatosis and fibrosis, the key histological parameters of NASH. Although our SHG-based microscopy techniques allow for objective quantitative assessment of steatosis and fibrosis changes on a continuous scale, there are some limitations to be addressed. First, in this study, the animals in the WD, WDF, and HFD groups lacked grade 2 steatosis, which may affect the steatosis analysis, and the collection of liver samples at shorter time intervals (such as every two weeks) might be helpful to cover the whole steatosis grade spectrum. Second, in our study, fibrosis stages 1-4 were observed in liver tissues from WDF+CCl4 and HFDF+CCl4 animals, whereas only fibrosis stages 0–2 were covered by WD, WDF, HFD, and HFDF groups. It has been reported that without the acceleration from CCl4, milder liver fibrosis may only be triggered by a high-fat or high-fructose diet over a prolonged period (Berardo et al., 2020; Kanuri and Bergheim, 2013; Reimer et al., 2020).

In conclusion, quantitative assessment using stain-free SHG/TPEF technology could better differentiate fibrosis and steatosis progression in mouse models of NAFLD. Although further studies are required, we found that qFibrosis co-localised with macrosteatosis could be useful for the sensitive and specific monitoring of liver fibrosis changes in NAFLD mouse models. These results suggest that, in future studies, SHG/TPEF technology should be considered to improve the accuracy of histological assessment in NAFLD preclinical models.

## Materials and methods

### Animals and diets

Male C57BL/6 mice were purchased from Beijing Vital River Laboratory Animal Technology Co. Ltd., China. They were maintained at 25°C with a 12 h light/dark cycle and allowed standard chow and water ad libitum until the time of the study. All protocols used in this study were approved by the Animal Experimental Ethical Committee of the Peking University People’s Hospital (No.2019PHC004). WD (42% kcal fat, 42.7% kcal carbohydrate; WD; TD120528) and 45% HFD (45% kcal fat, 35% kcal carbohydrate; HFD; MD12032) were purchased from Medicience Ltd. (Jiangsu, China), and standard chow (18% kcal fat, 58% kcal carbohydrate) was provided by SPF Biotechnology Co., Ltd (Beijing, China).

### Mouse models of NAFLD

Eight-week-old male C57BL/6 mice were randomly divided into six groups: (1) WD group; (2) WDF group, WD with a high sugar solution (23.1 g/L d-fructose and 18.9 g/L d-glucose) in drinking water; (3) WDF + CCl4 group, WDF plus intraperitoneal injection of CCl4 (10%, 2.5 μL/g of body weight), once a week; (4) HFD group; (5) HFDF group, HFD supplemented with a high sugar solution (23.1 g/L d-fructose and 18.9 g/L d-glucose) in drinking water; and (6) HFDF + CCl4 group, HFDF plus intraperitoneal injection of CCl4 (10%, 2.5 μL/g of body weight) once a week. Mice remained on the experimental diets until euthanised at the indicated times (0, 4, 8, 12, 16, 24, and 32 weeks on diet) to obtain specimens of different steatosis grades and fibrosis stages (n>3 per stage or grade).

### Liver histology analysis

The left lobe of the liver was paraffin-embedded, cut serially into 4-μm sections for direct SHG/TPEF imaging, and stained with H&E and picrosirius red staining, as previously described (Liu et al., 2017). Slides were evaluated by a blinded expert liver pathologist for histologic features, including grades for steatosis and stages of fibrosis, according to the NASH CRN scoring system (Kleiner et al., 2005). In addition, the CPA, as determined by the % area stained with Sirius Red, was quantified from the histological images using ImageJ (National Institutes of Health, Bethesda, MD, USA) as per our standard procedures.

### Image acquisition system

All images of the unstained tissue samples were acquired using a Genesis (HistoIndex Pte. Ltd., Singapore) SHG/TPEF imaging system. SHG microscopy was used to visualize collagen and steatosis structures were visualized using TPEF microscopy. The samples were excited with a 780 nm two-photon laser, the SHG signals were recorded at 390 nm, and the TPEF signals were recorded at 550 nm. Image tiles were acquired for each whole liver sample at 20× magnification with a 512 × 512-pixel resolution per tile, and each tile had an actual dimension of 200 × 200 μm^2^. Multiple adjacent image tiles were captured to encompass the entire tissue area on each slide.

### Establishment of qSteatosis and qFibrosis

Using the NASH CRN scoring system as the reference standard, automated measures of fibrosis and steatosis were developed, termed qSteatosis and qFibrosis, respectively. The sequential procedure for establishing the two indices comprised: (1) detection of fat vacuoles and collagen in different regions of the lobules, (2) quantification of defined architectural parameters that represented the characteristics of various histopathological features, (3) feature selection from the most correlated parameters, and (4) model construction, combining feature parameters into a single “signature” index for each of the two NASH histological components.

### Feature Detection and Image Quantification

#### Steatosis

To evaluate the severity of steatosis, fat vacuole candidates were detected as black holes in the TPEF channel and then further identified as either macro- or micro fat vacuoles using the Classification and Regression Trees method (Breiman et al., 1984) by examining the hole shape, cell structure, surrounding collagen, etc. The regions of steatosis consisted of not only fat vacuoles with high densities but also the hepatocytes around the vacuoles, which can be identified by image dilation in the binary image of fat vacuole detection. Similar to fibrosis quantification, dozens of steatosis parameters were extracted from the whole tissue image as well as the central vein (CV), portal tract (PT), and perisinusoidal (PS) regions to assess correlations with NASH CRN steatosis grades (Liu et al., 2020).

#### Fibrosis

To evaluate the severity of liver fibrosis, collagen was detected in the SHG channels within the tissue area. Using Otsu’s (1979) automatic threshold method, the collagen signal was identified from background noise. Then, multiple feature parameters were extracted from the collagen signals, such as length, width, density, and intersections. As liver tissue can be divided into three histological regions, namely CV, PT, and PS, feature detection was also performed in each region. Together with the overall quantification, approximately 100 fibrosis parameters (Chang et al., 2018; Liu et al., 2020) were extracted from each image to evaluate correlations with NASH CRN fibrosis stages.

#### Co-localisation

To evaluate the severity of fibrosis co-localised with steatosis, we evaluated fibrosis changes in the area close to the fat vacuoles in the perisinusoidal space. As steatosis was classified as macrosteatosis and microsteatosis, sub-analysis of fibrosis co-localised with each steatosis type was also performed by correlating co-localised fibrosis parameters with histological scoring.

#### Feature selection and model construction

For each animal model, fibrosis and steatosis parameters were correlated with the NASH CRN fibrosis stages and steatosis grades. The shared parameters, whose correlation ranked relatively high among these parameters in all animal models, were found with respect to fibrosis stages, steatosis grades, and time points for modelling. Linear models of these parameters were then created to identify differences between animals with different NASH scores.

#### Statistical Analysis

The AUC was used to illustrate the performance of the parameters and the linear model as its discrimination threshold varied. The advantages of using qFibrosis were illustrated by comparing the AUC values using CPA only. The Spearman correlation test was used to correlate the parameters with the CRN scores, and correlation coefficients were used to rank the parameters. The statistical significance level was set at *P*<0.05.

## Competing interests

Q.Y. and X.T. are employees of HistoIndex or its subsidiary, X.T holds stock options in HistoIndex. The other authors have no conflict of interest to disclose.

## Funding

This work was supported by grants from National Natural Science Foundation of China (NSFC) (82170584 and 81870406), China National Science and Technology Major Project for Infectious Diseases Control during the 13th Five-Year Plan Period (2018ZX09201002-001-005)

## Data availability

The datasets used and/or analyzed during the current study are available from the corresponding author on reasonable request.

## Author contributions

FL, XXW, XT, and HYR designed the study. XXW, JR, and XHL performed the experiments and supported the mouse experiments. FFL, NW, and HYR provided essential support and data interpretation. FL, XXW, QY, and XT drafted the manuscript. All the authors have read and confirmed the final version of the manuscript.

## Ethics approval

All protocols used in this study were approved by the Animal Experimental Ethical Committee of the Peking University People’s Hospital (No.2019PHC004).

## Notes

### Competing Interest Statement

The authors have declared no competing interest.

## References

Berardo, C., Di Pasqua, L.G., Cagna, M., Richelmi, P., Vairetti, M. and Ferrigno, A. (2020). Nonalcoholic fatty liver disease and non-alcoholic steatohepatitis: Current issues and future perspectives in preclinical and clinical research. Int. J. Mol. Sci. 21,.

Breiman, L., Friedman, J., Stone, C.J. and Olshen, R.A. (1984). Classification and Regression Trees. Philadelphia, PA: Chapman and Hall/CRC.

Brunt, E.M., Janney, C.G., Di Bisceglie, A.M., Neuschwander-Tetri, B.A. and Bacon, B.R. (1999). Nonalcoholic steatohepatitis: a proposal for grading and staging the histological lesions. Am. J. Gastroent. 94, 2467–2474.

Chalasani, N., Wilson, L., Kleiner, D.E., Cummings, O.W., Brunt, E.M., Unalp, A. and NASH Clinical Research Network. (2008). Relationship of steatosis grade and zonal location to histological features of steatohepatitis in adult patients with non-alcoholic fatty liver disease. J. Hepatolog. 48, 829–834.

Chang, P.E., Goh, G.B.B., Leow, W.Q., Shen, L., Lim, K.H. and Tan, C.K. (2018). Second harmonic generation microscopy provides accurate automated staging of liver fibrosis in patients with non-alcoholic fatty liver disease. PloS one. 13, e0199166.

De Rudder, M., Bouzin, C., Nachit, M., Louvegny, H., Vande Velde, G., Julé, Y. and Leclercq, I.A. (2020). Automated computerized image analysis for the user-independent evaluation of disease severity in preclinical models of NAFLD/NASH. Lab. Investig.; J. Tech. Meth. Patholog. 100, 147–160.

Diehl, A.M. and Day, C. (2017). Cause, pathogenesis, and treatment of nonalcoholic steatohepatitis. New Eng. J. Med. 377, 2063–2072.

Ding, Z.-M., Xiao, Y., Wu, X., Zou, H., Yang, S., Shen, Y., Xu, J., Workman, H.C., Usborne, A.L. and Hua, H. (2018). Progression and regression of hepatic lesions in a mouse model of NASH induced by dietary intervention and its implications in pharmacotherapy. Front. Pharmacol. 9.

Dulai, P.S., Singh, S., Patel, J., Soni, M., Prokop, L.J., Younossi, Z., Sebastiani, G., Ekstedt, M., Hagstrom, H., Nasr, P., et al. (2017). Increased risk of mortality by fibrosis stage in nonalcoholic fatty liver disease: Systematic review and meta-analysis: Dulai et al. Hepatol. 65, 1557–1565.

Ferguson, D. and Finck, B.N. (2021). Emerging therapeutic approaches for the treatment of NAFLD and type 2 diabetes mellitus. Nat. Rev. Endocrinol. 484–495.

Friedman, S.L., Neuschwander-Tetri, B.A., Rinella, M. and Sanyal, A.J. (2018). Mechanisms of NAFLD development and therapeutic strategies. Nat. Med. 24, 908–922.

Ganz, M., Csak, T. and Szabo, G. (2014). High fat diet feeding results in gender specific steatohepatitis and inflammasome activation. WJ Gastroenterol. 20, 8525–8534.

Ge, F., Lobdell, H., 4th, Zhou, S., Hu, C. and Berk, P.D. (2010). Digital analysis of hepatic sections in mice accurately quantitates triglycerides and selected properties of lipid droplets. Exp. Biol. Med. 235, 1282–1286.

Golabi, P., Paik, J.M., AlQahtani, S., Younossi, Y., Tuncer, G. and Younossi, Z.M. (2021). Burden of non-alcoholic fatty liver disease in Asia, the Middle East and North Africa: Data from Global Burden of Disease 2009-2019. J. Hepatol. 75, 795–809.

Hagström, H., Nasr, P., Ekstedt, M., Hammar, U., Stål, P., Hultcrantz, R. and Kechagias, S. (2017). Fibrosis stage but not NASH predicts mortality and time to development of severe liver disease in biopsy-proven NAFLD. J. Hepatol. 67, 1265–1273.

Huang, D.Q., El-Serag, H.B. and Loomba, R. (2021). Global epidemiology of NAFLD-related HCC: trends, predictions, risk factors and prevention. Nat. Rev. Gastroenterol. Hepatol. 18, 223–238.

Ipsen, D.H., Lykkesfeldt, J. and Tveden-Nyborg, P. (2020). Animal models of fibrosis in nonalcoholic steatohepatitis: Do they reflect human disease? Adv. Nutr 11, 1696–1711.

Kanuri, G. and Bergheim, I. (2013). In vitro and in vivo models of non-alcoholic fatty liver disease (NAFLD). Int. J. Mol. Sci. 14, 11963–11980.

Kleiner, D.E., Brunt, E.M., Van Natta, M., Behling, C., Contos, M.J., Cummings, O.W., Ferrell, L.D., Liu, Y.-C., Torbenson, M.S., Unalp-Arida, A., et al. (2005). Design and validation of a histological scoring system for nonalcoholic fatty liver disease. Hepatol. 41, 1313–1321.

Kleiner, D.E., Brunt, E.M., Wilson, L.A., Behling, C., Guy, C., Contos, M., Cummings, O., Yeh, M., Gill, R., Chalasani, N., et al. (2019). Association of histologic disease activity with progression of nonalcoholic fatty liver disease. JAMA Netw. 2, e1912565.

Liu, F., Chen, L., Rao, H.-Y., Teng, X., Ren, Y.-Y., Lu, Y.-Q., Zhang, W., Wu, N., Liu, F.-F. and Wei, L. (2017). Automated evaluation of liver fibrosis in thioacetamide, carbon tetrachloride, and bile duct ligation rodent models using second-harmonic generation/two-photon excited fluorescence microscopy. Lab. Investig.; J. Tech. Meth. Patholog. 97, 84–92.

Liu, F., Goh, G.B.-B., Tiniakos, D., Wee, A., Leow, W.-Q., Zhao, J.-M., Rao, H.-Y., Wang, X.-X., Wang, Q., et al. (2020). QFIBS: An automated technique for quantitative evaluation of fibrosis, inflammation, ballooning, and steatosis in patients with nonalcoholic steatohepatitis. Hepatol. 71, 1953–1966.

Martin, S. and Parton, R.G. (2006). Lipid droplets: a unified view of a dynamic organelle. Nat. Rev. Mol. Cell Biol. 373–378.

McPherson, S., Hardy, T., Henderson, E., Burt, A.D., Day, C.P. and Anstee, Q.M. (2015). Evidence of NAFLD progression from steatosis to fibrosing-steatohepatitis using paired biopsies: implications for prognosis and clinical management. J. Hepatol. 62, 1148–1155.

Nevzorova, Y.A., Boyer-Diaz, Z., Cubero, F.J. and Gracia-Sancho, J. (2020). Animal models for liver disease -A practical approach for translational research. J. Hepatol. 73, 423–440.

Otsu, N. (1979). A threshold selection method from gray-level histograms. IEEE Trans. Syst. Man Cybern. 9, 62–66.

Paik, J.M., Golabi, P., Younossi, Y., Mishra, A. and Younossi, Z.M. (2020). Changes in the Global Burden of chronic liver diseases from 2012 to 2017: The growing impact of NAFLD. Hepatol. 72, 1605–1616.

Ramot, Y., Zandani, G., Madar, Z., Deshmukh, S. and Nyska, A. (2020). Utilization of a deep learning algorithm for microscope-based fatty vacuole quantification in a fatty liver model in mice. Toxicol. Pathol. 48, 702–707.

Reimer, K.C., Wree, A., Roderburg, C. and Tacke, F. (2020). New drugs for NAFLD: lessons from basic models to the clinic. Hepatol. Int., 14, 8–23.

Roth, J.D., Feigh, M., Veidal, S.S., Fensholdt, L.K., Rigbolt, K.T., Hansen, H.H., Chen, L.C., Petitjean, M., Friley, W., Vrang, N., et al. (2018). INT-767 improves histopathological features in a diet-induced ob/ob mouse model of biopsy-confirmed non-alcoholic steatohepatitis. WJ Gastroenterol. 24, 195– 210.

Sethunath, D., Morusu, S., Tuceryan, M., Cummings, O.W., Zhang, H., Yin, X.-M., Vanderbeck, S., Chalasani, N. and Gawrieh, S. (2018). Automated assessment of steatosis in murine fatty liver. PloS one. 13, e0197242.

Sun, W., Chang, S., Tai, D.C.S., Tan, N., Xiao, G., Tang, H. and Yu, H. (2008). Nonlinear optical microscopy: use of second harmonic generation and two-photon microscopy for automated quantitative liver fibrosis studies. J. Biomed. Optics. 13,.

Tandra, S., Yeh, M.M., Brunt, E.M., Vuppalanchi, R., Cummings, O.W., Ünalp-Arida, A., Wilson, L.A. and Chalasani, N. (2011). Presence and significance of microvesicular steatosis in nonalcoholic fatty liver disease. J. Hepatol. 55, 654–659.

Terrault, N.A. and Pageaux, G.-P. (2018). A changing landscape of liver transplantation: King HCV is dethroned, ALD and NAFLD take over! J. Hepatol. 69, 767–768.

Tilg, H., Adolph, T.E. and Moschen, A.R. (2021). Multiple parallel hits hypothesis in nonalcoholic fatty liver disease: Revisited after a decade. Hepatol. 73, 833–842.

Vilar-Gomez, E., Calzadilla-Bertot, L., Wai-Sun Wong, V., Castellanos, M., Aller-de la Fuente, R., Metwally, M., Eslam, M., Gonzalez-Fabian, L., Alvarez-Quiñones Sanz, M., Conde-Martin, A.F., et al. (2018). Fibrosis severity as a determinant of cause-specific mortality in patients with advanced nonalcoholic fatty liver disease: A multi-national cohort study. Gastroenterol.155, 443–457.e17.

Wang, Y., Vincent, R., Yang, J., Asgharpour, A., Liang, X., Idowu, M.O., Contos, M.J., Daitya, K., Siddiqui, M.S., Mirshahi, F. et al. (2017). Dual-photon microscopy-based quantitation of fibrosis-related parameters (q-FP) to model disease progression in steatohepatitis. Hepatol. 65, 1891–1903.

Younossi, Z.M., Stepanova, M., Ong, J., Trimble, G., AlQahtani, S., Younossi, I., Ahmed, A., Racila, A. and Henry, L. (2021). Nonalcoholic steatohepatitis is the most rapidly increasing indication for liver transplantation in the United States. Clinical gastroenterology and hepatology: the official clinical practice journal of the American Gastroenterol. Assoc. 19, 580–589.e5.

